# A Careful Consideration of the Influence of Structure, Partial charges and Basis Sets on Collision Cross Sections of Monosaccharides when Comparing Values from DFT Calculated Conformers to those Obtained Experimentally

**DOI:** 10.1101/162305

**Authors:** Lukasz G. Migas, Christopher J. Gray, Sabine L. Flitsch, Perdita E. Barran

## Abstract

Molecular modelling is routinely employed to assign 3D structures to collision cross sections (CCSs) derived from ion mobility mass spectrometry experiments (IM-MS). The assignment of model structures to the experimental CCSs remains an ambiguous task, where one of several methods may be used to obtain a CCS from a given set of coordinates. The most reliable of the commonly used techniques, the Trajectory Method, starts with atomic coordinates which can be accompanied by partial atomic charges, obtained using *ab initio* methods. Here, we use lithiated α- and β-glucose ions as exemplar molecules to detect the effect conformational modification and changes to the partial charge distribution have on computed collision cross sections. Six popular charge schemes (Mulliken, APT, CHelpG, MK, HLY and NPA) were examined in combination with three functionals (Hartree-Fock, B3LYP and M05) and five basis sets (STO-3G, 3-21G, 6-31G, 6-31+G and 6-31G^*^) on twenty unique structures. Our findings indicate that molecular conformation makes a significant contribution to fluctuations of partial charges in Electrostatic Potential (ESP) and Mulliken charge scheme; Partial charges derived using Natural Population Analysis (NPA) and ESP methods are largely independent of functional and basis set selection; and both selection of the charge scheme and functional/basis set combination play a large role in the resultant CCS, often causing few percent fluctuations in the computed values.

## Introduction

Ion mobility-mass spectrometry (IM-MS) is an analytical technique that permits separation of gas-phase ions based on their molecular weight, charge and shape. The structural features of each ion are determined by measuring their arrival time distribution after traversing through an inert buffer gas under the influence of a weak electric field; drift-tube IM-MS measurements can directly determine the ions mobility, *K*,^1^ and hence the rotationally averaged collision cross section (Ω, CCS) *via* the Mason-Schamp equation, whereas travelling wave IM-MS (TWIMS-MS) determines the collision cross section (^TW^CCS) *via* calibration procedures.^2^ Both IM-MS techniques have been applied to a broad scope of species to elucidate structural information, including small drug-like molecules,^3,4^ carbohydrates,^5,6^ DNA and RNA,^7^ proteins^8,9^ and protein complexes.^9^ It has also used to study protein unfolding,^8^ refolding,^9,10^ proton and electron transfer mechanisms^11,12^ and solvent adduct effects on the protein structure.^13^ A great advantage of IM-MS is the ability to elucidate detailed structural information about the studied system by separation of isomers,^3^ protomers^14^ and conformers.^5^

The popularity of IM-MS has grown rapidly since the release of commercial instrumentation from Waters (Synapt and Vion instruments)^15^ and Agilent (qToF 6560); historically, IM-MS was carried out on in-house designed instruments with home-built drift tubes commonly filled with helium. Experimental measurements are often coupled with predictions for likely geometries, hence accurate algorithms to compute CCS from input coordinates are essential for the task. Due to the prevalence of experimental data, historically most algorithms utilised monoatomic nobel gases^16–19^ to compute theoretical CCSs whereas commercial IM-MS instruments typically use nitrogen. The number of algorithms to derive theoretical CCSs has boomed in recent years, with new methods appearing nearly every year.^20,21^ Significant progress has been made in order to create accurate and efficient algorithms that utilise polyatomic gases (i.e. N_2_ or CO_2_), in particular by Campuzano *et al*.,^22^ Bleiholder *et al*.^23^ and Larriba *et al.*^24^ Since the majority of experimental data to date has been reported as helium CCSs, a common practice amongst the community using the TWIMS instruments is to represent the nitrogen measured arrival time distributions as effective helium collision cross sections and therefore take advantage of computational methods that use helium as the buffer gas. This methodology has been first developed by Ruotolo^2,25^ and subsequent studies by Bush *et al*.^26^ and Shvartsburg *et al*.^27^ have demonstrated that this approach produces only minor errors (<3 %) for protein complexes and even lower typical errors for small molecules, depending on the selection of calibrants and the quality of the calibration procedure.

A number of recent studies have utilised high-level DFT calculations to optimise potential candidate models to correlate experimental and theoretical results. For instance, Warnke *et al.*^14^ demonstrated that combining IM-MS and infrared multiple photon dissociation (IRMPD) spectroscopy reveals two distinct protomers of benzocaine, where depending on the protonation site (N or O), a unique N-H and O-H stretch frequencies appear. Similarly, Voronina *et al*.^28^ utilised IM-MS and cold-ion spectroscopy to study the bradykinin fragment (BK[1-5]^2+^) ions elucidating *cis-trans* and *trans-cis* conformations for the more compact conformer and kinetically trapped *trans-trans* configuration for the larger conformer, highlighting the *cis-trans* isomerization of the peptide bonds during the desolvation process. In both cases, IM-MS data was supplemented with gas-phase IR experiments which in turn were supported by high-level calculations to assign potential *in vacuo* species. Recent work by Boschmans *et al*.^29^ combined ^TW^CCS measurements and theoretical CCS calculations of several small molecules which were observed to have multiple conformers. To propose 3D model structures they utilised high-level density functional theory (DFT) optimisation of multiple conformers using two different levels of theory; each resulted in marginally different CCSs due to the minor changes to the structure introduced during the optimisation process (namely small adjustments to bond lengths, bond angles and the dihedral angles). In another study, Young *et al*.^30^ focused on determining the effect analyte charge distribution has on CCSs of carbon clusters ranging from C_24_ to C_960_ and charge states of +1 to +6. While the atomic charges were not evaluated using DFT or *ab initio* methods, their findings highlight the importance of charge distribution on the CCS, in particular for gases with high polarizability volume (*i.e.* N_2_).

Computation of theoretical CCSs requires the 3D atomic coordinates and corresponding partial charge distribution to accurately determine ion-neutral collision interactions; partial charges are more important for nitrogen calculations, since its polarizability volume is approximately ten times that of helium (*α’* _N2_=1.7 Å^3^, *α’*_He_=0.21 Å^3^) resulting in a much deeper ion-neutral potential well. Atomic partial charges can be calculated using multitude of methods, most commonly Mulliken population analysis,^31^ electrostatic potential methods,^32–34^ Natural Population Analysis^35^ along with several others.^36–39^ Each different charge scheme method yields unique atomic charge distributions, often dependent on the size of the basis set, implying a significant impact on the resultant collision cross sections of the ions, in particular for species with high surface charge density where the ion-neutral interactions would be most prevalent and when using polar buffer gases.

In this work we focus on singly charged, lithiated α- and β-disaccharide fragment ions (C-ions) which we have shown (Gray *et al*.^5^) to retain an anomeric configuration during the glycosidic bond fragmentation and exist as two separable species in the IM-MS experiment, as shown in Figure 1a. The discovery of the ‘memory effect’ has significant implications for an rapid MS based method to determine the stereochemistry of glycans, previously mostly achieved by combinations of low throughput NMR, glycosidase digestions coupled with chromatography and chemical derivatisation and tandem MS.^40^

**Figure 1.**
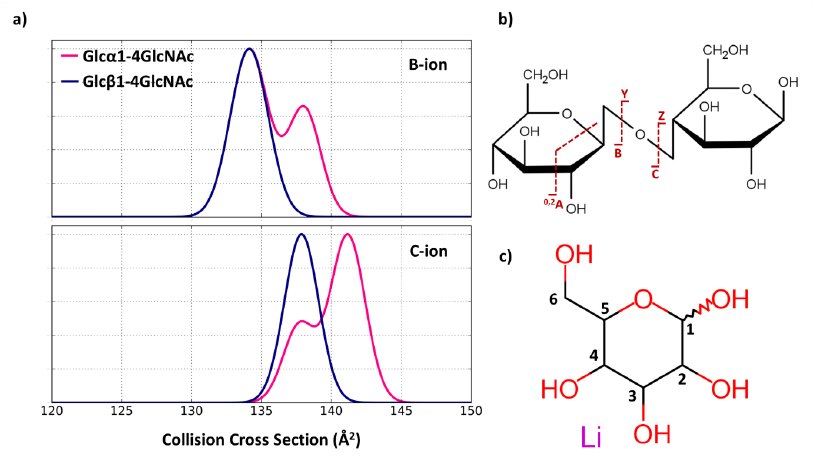
a) Travelling-wave ion mobility mass spectrometry collision cross section distributions of Glcα1-4GlcNAc and Glcβ1-4GlcNAc fragment ions obtained from tandem MS experiments. Product species generated by CID of the α-precursor (magenta) result in two species for both B- and C-fragment ions, whereas β-product ions (blue) consist of single species; Data shown in this figure has been adapted from Figure 2 from Gray *et al*.^5^ here, reporting Glcα1-4GlcNAc and Glcβ1-4GlcNAc only. b) Scheme representing the Domon-Costello nomenclature for carbohydrate fragmentation^47^; c) Carbon labelled monosaccharide showcasing the numbering system.

Our study seeks to highlight and answer three primary questions; ‘How much does the molecular conformation affect the partial charge distribution?’, ‘How much do commonly employed charge schemes depend on selection of the functional and the size of the basis set?’ and ‘Does the partial charge distribution affect the theoretical CCSs of the same ions?’

We demonstrate that in the case of simple monosaccharides, theoretical CCSs are heavily dependent on the selection of the charge scheme, basis set and in some cases the functional. The variation based on chosen charge scheme and basis set in the partial charge distribution on structurally identical ions can lead to as much as 3-8 % change in the calculated CCS. This indicates that the approach to matching experimental CCS values with those obtained from calculated structures requires considerable care.

## Experimental and Computational Methods

### Travelling wave ion mobility-mass spectrometry

Samples were prepared according to the conditions described previously.^5^ Carbohydrates were diluted to 5 pmol/μL with 1:1 methanol:water prior to analysis; to study lithiated glycans, 10 eq. of lithium chloride was doped into the solution. Samples were infused into a Synapt G2 HDMS (Waters, UK) by nanoelectrospray ionization using in-house pulled borosilicate tips (World Precision Instruments, USA, thin-wall capillary, 4” length, 1.2 mm O.D). The capillary, cone voltage and source temperature were typically set to 0.8-1.5 kV, 40 V and 80 °C respectively. The IM travelling wave speed was set to 1200 m/s and wave height set to 40 V. Nitrogen drift gas flow was set to 90 mL/min for all experiments. The helium and argon flow was set to 180 and 2 mL/min respectively for the helium and trap cell. CID of quadrupole selected precursor ions was induced in the trap region using an argon collision gas at 30 V. Data were collected between *m/z* 50-1200 with a product ion tolerance of ± 50 ppm. Mass measurements were calibrated using product ions following CID of Glu-fibrinopeptide (trap CE 35 V) by infusion (500 fmol/μL, 1:1 acetonitrile:water, 0.1 % (v/v) formic acid) at a flow rate of 0.1 μL/min. Drift times were calibrated to calculate ^TW^CCS_N2_ values by infusion of a stoichiometric mixture (1 pmol/μL, 1:1 acetonitrile:water, 0.1 % formic acid) of small molecules (acetaminophen, verapamil, *N-*ethylaniline, colchicine and reserpine; all Sigma), whose CCS have been previously verified.^22,41^ Mass spectra and ATDs were recorded in triplicate and processed using MassLynx v4.1 (Waters, UK) and OriginPro 9.1 (OriginLabs, USA) respectively. ATDs were calibrated and subsequently normalised to their maximum intensity. Gaussian distributions were fitted to these spectra and the centre of the fitted peak was taken as the peak CCS.

### Conformer generation

Conformational searches were carried out using the *sander* module in Amber14^42^ with the GLYCAM_06j forcefield.^43^ Molecular dynamics was used to explore conformational landscape of the α- and β-Glc ions and the position of the Li^+^ cation adduct. Each molecular system was heated to 4000 K and dynamics were performed for 20 ns, using a time step of 1 fs to ensure energy conservation. Evaporation of the Li^+^ cation was prevented by adding distance restraint around the molecule; a restraint of 10 Å with force constants of 0 and 20 kcal/mol (lower and upper boundaries) was found sufficient to prevent the loss of the adduct ion. Structures were extracted every 200 steps (0.0002 ns, 100,000 models) and filtered based on their similarity (root square mean deviation, RMSD) and energy of the system; duplicate models were removed, significantly reducing the number of candidates. Remaining models were optimised using a semi-empirical PM6-DH+ in MOPAC2016;^44,45^ optimised models were subsequently filtered based on their RMSD and energy, leaving a small number of candidate ions. Remaining structures were refined using Density Functional Theory (DFT) in Gaussian09^46^ at the B3LYP/6-311+G^*^ level of theory. Resultant pool of structures was manually filtered to select structurally and conformationally varied molecules.

### Calculation of partial charges

Partial atomic charges were evaluated from a single-point calculation with either Hartree-Fock (HF) or density functional theory (DFT) B3LYP or M05 functional. Five Pople basis sets were included: STO-3G, 3-21G, 6-31G, 6-31+G and 6-31G^*^. Six charge schemes were considered in the calculations, namely Mulliken,^31^ Atomic Polar Tensor (APT),^39^ Natural Population Analysis (NPA),^35^ Merz-Singh-Kollman (MK),^32^ CHelpG^33^ and Hu-Lu-Yang (HLY).^34^ The Mulliken charge scheme is historically the most important method to derive partial charges. It relies on calculating charges from the atomic orbitals on the wavefunction of the molecule; one known deficiency of the Mulliken charge scheme is its dependence on basis sets. The NPA method offers a more refined wavefunction-based method that calculates charges based on natural atomic orbitals giving more stable charges with varied sizes of the basis set. The electrostatic potential (ESP) methods are calculated by trying to reproduce the molecular electrostatic potential using a large number of grid points-the methods differ in the way the grid points are selected. The CHelpG scheme places points in a cubic box that encompasses the molecule whereas the Merz-Kollman method uses 4 layers encompassing the molecule. Each layer is scaled by a scaling factor to result in surface larger than the Van der Waals surface. The HLY charge scheme utilises an object function of the entire molecular volume space instead of discreet grid points surrounding the molecule. The object function is used to prevent variations due to the molecular orientation and improve the numerical stability.

### Collision cross section calculations

Theoretical rotationally averaged CCSs were evaluated using MOBCAL_N2_ using the trajectory method only.^18,22^ The code was modified at line 265 to increase the number of points in Monte Carlo integration of the impact parameter (*imp*) from 1000 to 1500 to improve the convergence of the calculation and reduce standard deviation of the calculation; typical standard deviations were ~0.5 %, with all being less than 1.2 %. Gaussian09 output files (.log) were converted to the MOBCAL format (.mfj) using an in-house script allowing batch processing.

## Results and Discussions

### Ion mobility mass spectrometry of diglucosides and fragment ions

Figure 1a shows the CCS distribution of B- and C-fragment ions of lithiated Glcα/β1-4GlcNAc diglucosides generated by collision-induced dissociation (CID) prior to ion mobility separation (as shown in Figure 1b). The B- and C-product ions of α-linked diglucosides revealed two peaks in the IM-MS profiles centered at 134.1 and 137.9 Å^2^ for B-ions and 137.9 and 141.3 Å^2^, whereas a single peak was observed for the analogous β-linked structures with collision cross sections of 134.1 Å^2^ for B-ion and 137.9 Å^2^ for C-ion. Other saccharides and product ions including permethylated variations are discussed in greater detail in a separate manuscript.^5^

Although the difference in the measured collision cross sections of B- and C-fragment ions is < 4 Å^2^, comparison to high-level computational models can yield useful information with regards to the molecular structure and fragmentation mechanisms. Computation of theoretical CCSs relies on accurately determined 3D coordinates of the ion and the partial atomic charges, so the ion-neutral interactions with the buffer gas can be appropriately determined. Molecules with highly localised charge density (*i.e.* cation adduct) will be involved in stronger interactions with the buffer gas, than those with highly delocalised charge. In order to assess the sensitivity of the CCS calculations to the molecular structure and partial charge distribution changes, twenty unique structures of α- and β-ions were selected to represent the potential models of the C-ion data. The structural assignment of the B-ions is more challenging due to the high number of isomeric structures it can attain during the fragmentation process. Since we are more interested in quantifying the impact charge schemes and the selection of the basis set has on the calculated CCS, the studied system should be conformationally and not structurally complicated.

### Molecular conformations of α-/β-fragment ions

Even simple monosaccharides represent a challenging group of molecules for computational chemists due to their vast number of available conformations caused by the presence of five hydroxyl groups (for hexoses) and multitude of puckering configurations, and this is further complicated by the addition of adduct ions (*i.e.* Li^+^, Na^+^, Ca^2+^) that can coordinate in a number of ways with the carbohydrate. In order to determine the effect the conformation of the ion has on the partial charge distribution and CCSs, ten conformations for each α- and β-Glc ions were selected. Models were based on the glucopyranose structure (and not the linear form) and manually selected to give high variability in the orientation of the hydroxyl groups and position and coordination number of the cation but also to represent the likely puckering configurations, either ^1^C_4_ and ^4^C_1_. Rather than only selecting the lowest energy structures from the global/local minima, a number of more energetic conformers were chosen to diversify the group of candidates thus providing a better framework to study the conformational dependence of charge schemes. The free energy of the monosaccharides can differ by few kcal/mol simply due to slight differences in the orientation of the hydroxyl groups,^48^ leading to increased or decreased intramolecular hydrogen bonding network, hence identification of *true* global minima for even simple monosaccharides is a non-trivial task.

Following extensive conformational search, ions with the lowest energy were found to coordinate lithium cation with hydroxyl groups at positions OH3 and OH4. Structures of these are shown in Figure S1, denoted **βGlc(1)** and **αGlc(2)**. In solution, both α- and β-Glc prefer the ^4^C_1_ conformation, which were predominantly seen during the conformational search. Models in the ^1^C_4_ configuration were found to be higher in energy, in agreement with Mayes *et al.*^48^ The 3D models of α-Glc configurations are shown in Figure S1a, depicting multiple coordination sites of the lithium cation; conformers **αGlc(1-3,5,7-10)** are in the ^4^C_1_ conformation and **αGlc(4,6,8)** in the ^1^C_4_. Similarly, the β-Glc models are shown in Figure S1b, where all conformations were in the ^4^C_1_ configuration. Minor differences in the conformation of the ion will have an impact on the charge distribution across the molecule, which in turn would impact the ion-buffer interactions in the mobility experiment and theoretical CCS calculations. The boundary dividing the electron density between different atoms in a molecule has not been found in nature, hence the partitioning of total electron density across multiple atoms is utterly subjective and quite often basis set dependent. To examine and alleviate the ambiguity of partial charge division across the molecule, six popular charge schemes have been chosen to assign the partial charge distribution to each conformation, and determine their dependence on the molecular conformation and the basis set. We investigated Mulliken population analysis, Atomic Polar Tensor (APT), electrostatic potential derived charges (Merz-Singh-Kollman, MK; Hu-Lu-Yang, HLY and CHelpG) and Natural population analysis (NPA). The partial charge schemes differ in their method of assigning values to particular atoms, often resulting in a broad range of values for the same atoms; each method is constrained to represent the net charge of the ion of +1 in the case of α-/β-Glc. Any charge scheme should be sensitive to the chemical environment each atom experiences, in the case of lithiated monosaccharides the position of lithium should play a crucial role in the partial charge assignment.

### Impact of ion conformation and basis set selection on the partial charge distribution

To probe the impact of molecular conformation on the partial charge distribution and determine the dependence of the charge schemes on the selection of functional and basis set, each conformer of α- and β-Glc was subjected to single point calculation using either HF, B3LYP or M05 functional with STO- 3G/3-21G/6-31G/6-31+G or 6-31G* basis set and one of six charge schemes. The 3D configuration of each conformer remained intact between each calculation. Partial charges from all conformations (including both α- and β-Glc ions) are represented together to give average partial charges for each atomic centre; the width of the error bars represents the most extreme charge fluctuations caused by the change in the molecular structure. Large fluctuations in the average partial charges indicate significant dependence of that charge scheme with particular functional/basis set combination on the ion conformation. The average partial charges computed using Mulliken population analysis and APT charge schemes are shown in Figure 2a and b, respectively.

**Figure 2.**
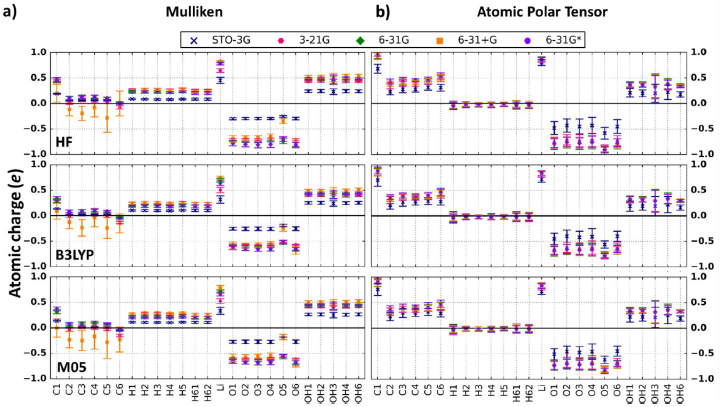
Average partial atomic charges from all conformers of α-/β-Glc ions determined using HF, B3LYP and M05 functionals with several basis sets: STO-3G (blue), 3-21G (magenta), 6-31G (green), 6-31+G (orange) and 6-31G^*^ (purple). a) Partial charges from Mulliken charge scheme; b) Partial charges from Atomic Polar Tensor. Average charges were computed from 20 conformers whereas the standard errors were evaluated by determining the maximum spread of charges on each atom centre due to the conformational differences. Average partial charges for MK, HLY, CHelpG and NPA can be found in SI Figure S2 and S3.

The Mulliken charge scheme was found to have a mild conformational dependence with typical charge fluctuations of <±0.08 e for each basis set. Charges computed with the 6-31+G (orange) basis set were significantly more varied on the carbon atoms (C1-C5, > ± 0.25 e), whilst the fluctuations on other atom types were unaffected, although the magnitude was altered to offset the more negative charges on the carbon atoms. Unsurprisingly, the partial charges on the lithium adduct varied more than the remaining atoms, regardless of the basis set, since Li is involved in the coordination to the hydroxyl groups in either mono-, bi- or tridentate modes. The broad dispersion of partial charges for the same atoms was caused by the changes to the size of the basis set indicating high dependence on the basis set. Significant fluctuations on the carbon atoms were largely caused by the augmented 6-31+G basis set, leading to more negative charges than for the remaining basis sets. The minimal basis set (STO-3G, blue) led to broadly different partial charges for carbon hydrogens (H1-H62; H^C^), hydroxyl oxygens (O1-O4, O6; O^OH^) and hydrogens (OH1-OH6; H^OH^), ring oxygen (O^COC^) and lithium adduct in comparison to the other basis sets. The choice of the functional played only a minor role in changing the magnitude of the charges, resulting in slightly more positive charges on the lithium adduct (by 0.05 *e*) with the HF functional than the B3LYP/M05. Ion conformation of α-/β-Glc had little impact on the aliphatic atoms when the partial charges were evaluated using the APT charge scheme; the fluctuations on carbon and H^C^ atoms was typically < 0.03 *e* for most basis sets. The charges on the hydroxyl atoms were more varied with fluctuations of ±0.11 *e* on O^OH^ and ±0.08 *e* on hydroxyl hydrogens H^OH^; interestingly, the variation on H^OH^3^^ atom which was frequently involved in coordination to Li increases to ±0.25 e. The smallest STO-3G basis set resulted in elevated charge fluctuations for all atomic types, whilst the remaining basis sets presented more subdued results. Neglecting the results obtained with STO-3G, all functional and basis set combinations resulted in narrow partial charge distribution (within < 0.03 *e* differences); the small dispersion of partial charges for each atomic centre with multiple basis sets indicates independence from molecular structure and basis sets.

Electrostatic potential (ESP) methods determine partial atomic charge by least-square fitting a number of points around each atom in the molecule to represent a molecular electrostatic potential (MEP); each ESP method uses different approaches to derive the atomic charge, primarily by changing the distance range at which the points are calculated. Unsurprisingly, charges calculated using different ESP methods for the same molecules (i.e. MK, CHelpG, HLY or many others) will be typically different. A number of studies had shown that altering the molecular conformation of the ion will result in fluctuations to the atomic charges, in particular for atoms with deeply buried atoms due to the deformations of the Van der Waals surfaces.^49^ Average partial charges from Merz-Kollman and Hu-Lu-Yang charge schemes are shown in Figure S2a and S2b, respectively, whilst CHelpG charges are shown in Figure S3a. In the case of α-/β-Glc monosaccharides, the molecular conformation was found to have significant impact on the partial charge distribution resulting in large variations on most atomic types. Partial charges on H^C^ and Li showed the smallest dependence on the molecular conformation for each ESP method (±0.07 *e* and ±0.05 *e*, respectively), whilst charges on carbon atoms were consistently high for each of the methods (±0.24, ±0.26 and ±0.23 *e* for MK, HLY and CHelpG methods, respectively). Similarly high fluctuations were observed for the hydroxyl groups (±0.15 *e* for O^OH^ and ±0.10 *e* for H^OH^), and ring oxygen (O^COC^, ±0.13 *e*). Importantly, the amount of variation was consistent between multiple functionals and basis set combinations, hence the large fluctuations can be solely attributed to the molecular conformation. Whilst the ESP methods showed pronounced sensitivity to the conformation of the molecule their dependence on the basis sets is more systematic, displaying better behaviour than Mulliken and akin to the APT method. Neglecting again the smallest basis set STO-3G, the remaining functional and basis set combinations give excellent agreement on all atomic centres. The fluctuations on lithium and hydroxyl groups were in particular very low, typically within ±0.03 *e* of one another. Despite differences in the fitting procedures, the ESP methods resulted in partial charges that were only marginally different from one another.

While the other charge schemes showed certain levels of susceptibility to the molecular conformation and the size of the basis set, the Natural Population Analysis method showed entirely different behaviour to the changes in the structure and level of theory (Figure S3b). The fluctuations caused by structural changes were typically < ±0.02 *e* on the hydrocarbon atoms (C and H^C^) and < ±0.05 *e* on O^COC^ and the hydroxyl atoms. A wider range of charges was observed for Li adduct (±0.07 *e*) and H^OH^3^^ (±0.08 *e*). Again, neglecting the minimal STO-3G basis set, the remaining basis set and functional combinations resulted in practically identical partial charges. Slight deviations of < ±0.04 *e* for Li, O^OH^ and O^COC^ was observed with the 3-21G basis set, however each of the 6-31G basis behaved remarkably well.

The minimal basis set STO-3G is inadequate for partial charge calculations, routinely resulting in drastically different partial charges for majority of the atomic types whereas larger basis sets, including the 3-21G were found to give smaller partial charge fluctuations. The Mulliken charge scheme showed mild fluctuations caused by the molecular conformation charges and strongest dependence on the basis set selection making it an inadequate source of partial charges. The APT and ESP methods displayed much better behaviour, obtaining less varied partial charge distributions for the majority of atomic types; taking the conformational dependence into account, the APT showed much lower fluctuations for certain atom types (H^C^, C and Li) than the ESP charge schemes. The NPA charge scheme was found to result in the least varied average partial atomic charges, exhibiting weakest dependence on the basis set selection, while also being least sensitive to molecular changes. It is no surprise that each of these methods results in partial charges of completely different magnitude. For comparison, Figure S4 shows the average partial charge from each charge scheme for each quantum mechanical method with 6-31G basis set (shown without the error bars for clarity purposes). The largest observable variations are shown for the carbon atoms charges varied by as much as 0.5 *e* (*e. g.* C1: APT = 0.98 *e* and NPA = 0.47 *e*). Charges on the hydrogen atoms (H^C^) were less varied, yet ranged between 0.3 *e* (NPA) to −0.04 *e* (APT) with the ESP methods in-between at 0.12-0.15 *e*. The agreement for Li was remarkably good, with fluctuations < 0.05 *e* between the methods, while the values on O^OH^ were in good agreement with partial charges of approx. −0.8±0.1 *e*. The charges on H^OH^ were in excellent agreement with the exception of APT which predicted values of ~0.38 *e* whilst all other charge schemes of ~0.52 *e*. Similarly, the charges on O^COC^ were closely related with the exception of values computed with APT method (APT = −0.9 *e* and other methods = −0.65±0.1 *e*).

### Impact of partial atomic charges on collision cross sections

As an example to demonstrate the partial charge fluctuations and their impact on CCS values on *static* carbohydrate ions, Figure 3a shows the heavy atom partial atomic charges computed using APT, MK and NPA charge schemes at M05/6-31G level of theory for βGlc(1) conformer. While the molecular structure remains constant, as demonstrated by the identical values of biophysical properties such as radius of gyration, dipole moment, surface area and volume, the collision cross section changes substantially (as shown in Figure 3b). The observed differences in the computed collision cross sections can be attributed to the fluctuations in the partial charge distribution. Individual atomic charges for the single βGlc(1) conformer are shown in Figure 3c; the global trends in partial charge fluctuations for each charge scheme were discussed earlier on, however focusing on a single example it is apparent that the choice of the charge scheme has substantial effect on the computed CCS values. In this case, the APT charge scheme which was shown to reduce the magnitude of partial charge on the H^C^ and H^OH^ atoms resulted in 1.57 % smaller CCS. In contrast, the NPA charge scheme increases the magnitude of the charge on the H^C^ atoms and leads to 2.72 % larger CCS. While this is an isolated example, discussion below showcases how similar trends are observed for all tested computation methods.

**Figure 3.**
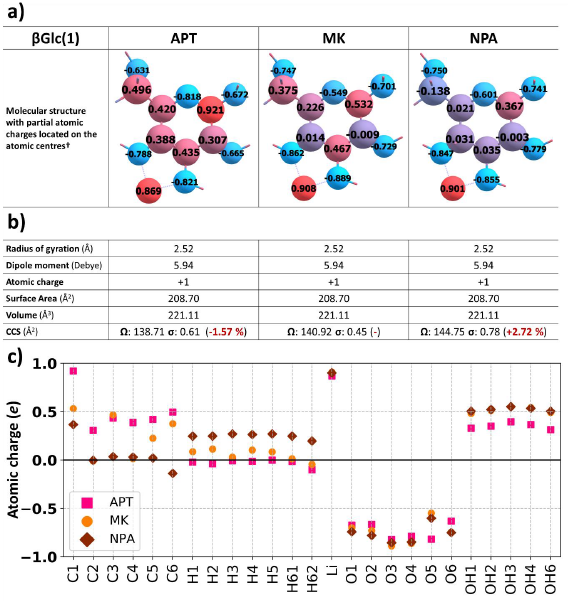
Partial charge distribution of heavy atoms on single conformer of βGlc(1) calculated using M05/6-31G basis set. a) the comparison between the APT, MK and NPA charge schemes reveal large variations in the magnitude of partial atomic charges on each atom resulting in significant fluctuations in the computed CCS values; other physical parameters are unaffected. Atoms with positive partial charge are shown in red whilst negatively charged atoms in blue; b) Overview of physical properties for each β Glc(1) molecule; c) Plot of partial charges on βGlc(1). Molecule visualised using Chemcraft 1.8.^50^ † Partial charges of hydrogens are not shown for clarity purposes.

CCSs computed with partial charges from Mulliken charge scheme gave a broad CCS distribution (Figure 4a), predominantly dominated by the significantly smaller values (> 5 Å^2^) caused by the large discrepancies in partial atomic charges computed with the minimal basis set STO-3G. Good agreement was observed for the CCS values obtained with 3-21G, 6-31G and 6-31G* basis sets, with CCS fluctuations of < 2 Å^2^. In contrast, the CCSs calculated with partial atomic charges from 6-31+G basis set were larger (by > 3 Å^2^ for most conformers). Small fluctuations caused by the selection of the quantum mechanical method were also observed indicating that even small changes in the partial atomic charges have direct impact on the computed CCSs. Differences of < 0.03 *e* on few atomic centres obtained with HF functional resulted in CCS 4 Å^2^ larger than those obtained for the same conformer with B3LYP/M05 functionals. The CCSs calculated with APT partial charges were in good agreement between different basis sets and functionals (Figure 4b). Akin to the Mulliken charge scheme, CCSs computed with STO-3G basis set were ~3-5 Å^2^ smaller for the other basis sets. Since the partial charges for most atoms were in good agreement, the CCSs are typically within the 0.5-1 % error boundary of the calculation. Altering the quantum mechanical method introduced no additional deviation to the CCS calculation, with all values falling within 1 Å^2^ for the three functionals. The ESP methods were found to have good agreement in the partial charge distributions for multiple functional and basis set combinations and their CCS values reflected this behaviour (Figure S5 and S6a). As with the APT and Mulliken methods, CCSs calculated with STO-3G partial charges were > 5 Å^2^ smaller for each conformer than with the other basis sets. The CCSs computed with 6-31+G basis set were on average marginally larger than the other methods, however the values were within the error margins of the TM calculation. The impact of the functional was relatively low, with minor CCS alterations for the larger basis sets; the CCSs calculated with HF were consistently larger than B3LYP/M05 values, however within the error of the calculation. Identical behaviour was exhibited by each of the ESP schemes, in good agreement with the partial charge observations. The NPA charge scheme (Figure S6b) showed the best agreement in partial atomic charges and calculated CCS using each functional and non-minimal basis set. The CCS agreement for values calculated using 6-31G, 6-31+G and 6-31G* basis was within 0.5-1 % error boundary of the calculation; similarly, the fluctuations in charges caused by the different quantum mechanical methods led to insignificant differences in the CCS values. As was the case for the other charge schemes, CCSs calculated with STO-3G and to some extent with 3-21G basis sets were underestimated by as much as 8-10 Å^2^ (STO-3G) and 2 Å^2^ (3-21G) for each conformer.

**Figure 4.**
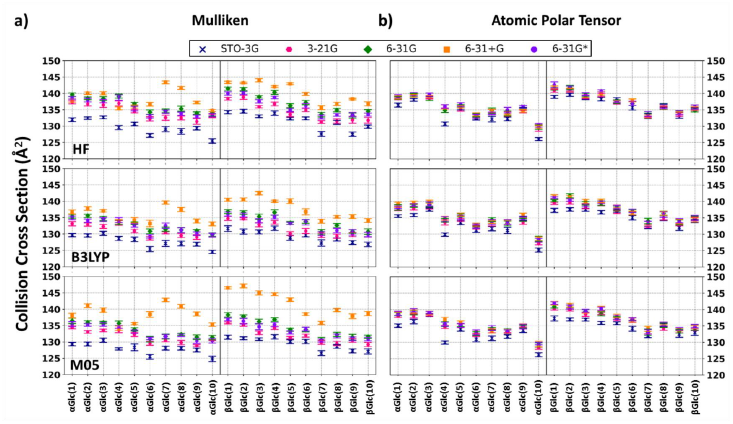
Rotationally averaged collision cross sections calculated using the Trajectory Method with partial charges calculated using HF, B3LYP or M05 functional with several basis sets: STO-3G (blue), 3-21G (magenta), 6-31G (green), 6-31+G (orange) and 6-31G* (purple) for a) Mulliken, and b) Atomic Polar Tensor charge schemes. Error bars on each data point represent the standard deviation from Trajectory Method calculation, typically less than 0.5 %, and always less than 1.2 %. CCS plots for MK, HLY, CHelpG and NPA can be found in SI Figure S5 and S6.

Although the majority of ion mobility studies employ the Merz-Kollman charge scheme (using the B3LYP functional and 6-31G or 6-311G basis sets), the results demonstrated herein clearly advocate for a more cautious approach to computing theoretical collision cross sections for small molecules. A wide number of charge schemes, functionals and basis sets are available, each potentially resulting in a unique partial charge distribution and consequently unique collision cross section, even for the same 3D structure. The collision cross section distribution (CCSD) as compared to the experimentally observed range is shown in Figure 5 for α-Glc (b) and β-Glc (c) species. In both types of calculations, the minimal basis set STO-3G resulted in dramatically different partial charges and considerably lower CCS. To some degree, the 3-21G basis set led to slight underestimation of the CCSs, in particular for the NPA charge scheme. CCSs computed with the 6-31G basis sets (including the polarizability and dispersion augmentations) occupied similar CCS distributions for each charge scheme, apart from NPA. Values computed with Mulliken and APT charge schemes tended to be smaller than the ESP methods, while the NPA charge scheme led to larger CCSs for the same conformers. The APT and NPA charge scheme resulted in broader CCSDs than ESP, surprisingly permitting clearer differentiation of the α and β-Glc conformers, despite displaying weaker dependence on molecular conformation in the partial charge calculations as shown in Figure 2, S2-3.

**Figure 5.**
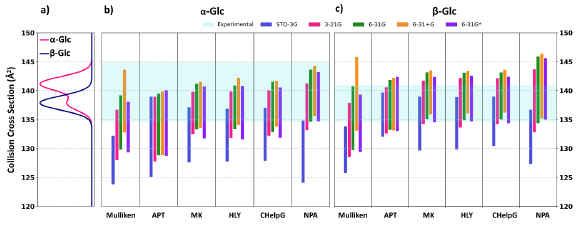
Collision cross section distribution (CCSD) generated by each partial charge scheme for all conformers of α-/β-Glc ions calculated for each basis sets with B3LYP functional. a) Experimental CCSD; b) CCSD of α-Glc ions; c) CCSD of β-Glc ions. The CCSDs were obtained by extracting the smallest and largest CCS value for each basis set (from all functionals) incorporating the average standard error for the conformer ensemble of α-/β-Glc ions. The cyan overlay represents the width of the experimental Gaussian distribution (shown in Figure 1a); horizontal black lines represent the apex values for the two observed conformers of α-Glc and one for β-Glc ion.

In many cases the IM-MS does not yield baseline separation of conformers/protomers, and rather results in similar arrival time distributions as in the case of α- and β-Glc fragment ions which are separated by <4 Å^2^ (Figure 1a). The results presented above suggest that unambiguous assignment of CCSs to structural models is more complicated than initially thought. The traditional route of *structure optimisation*, *calculation of partial atomic charges* and *computation of collision cross sections*, as often employed in many studies, does not account for the minor (and major) fluctuations in the partial charge distribution associated with different charge schemes and basis sets.

## Conclusions

In this report, we report the impact of partial atomic charges computed using a number of functionals, basis sets and charge schemes on 20 α-/β-Glc conformations. In answer to our initial questions, molecular conformation results in alterations to the partial charge distribution, in particular for the ESP methods (MK, CHelpG and HLY), whilst the other methods showed weaker dependence. Secondly, all charge schemes with the exception of Mulliken population analysis showed little dependence on the functional and basis set, with APT and NPA displaying nearly complete invariance. Lastly, our data indicates that molecular structure, partial charge distribution and resultant collision cross sections calculated using the Trajectory Method are strongly interlinked and small differences in the partial charges, whether caused by the conformational or quantum chemical method have significant impact on the computed CCSs. Overall, partial atomic charges of α-/β-Glc ions computed with NPA and ESP charge scheme result in reliable and predominantly functional and basis set independent charges, assuming minimal basis sets are omitted. Each of the examined ESP methods showed great sensitivity to conformational changes and little basis set dependence, making them a good candidate for reliable partial charge derivation for CCS calculations; careful considerations should be taken for large systems, for which the ESP method is known to inaccurately determine partial charges. In those cases, we advise that the NPA charge scheme should be used.

## Data availability

The interactive IPython (Jupyter) notebooks containing experimental and computational results from this study are available to view on GitHub at https://github.com/BarranLab/ChargePaper_2017.

## Acknowledgements

We gratefully acknowledge BBSRC and the Waters Corporation for support of this work in the award of a DTP studentship to L.G.M. We also thank the British Mass Spectrometry Society (BMSS) for funding our nanospray tip puller and University of Manchester for continued support of our research.

## Author contributions

With initial direction from P.E.B and S.L.F., L.G.M initiated and carried out all computational aspects of the study. L.G.M and C.J.G. carried out the mass spectrometry experiments. L.G.M. analysed the data and drafted the manuscript. All authors contributed towards final editing of the manuscript.

